# Virulence of *Fusarium oxysporum* strains causing corneal or plant disease is associated with their distinct accessory chromosomes

**DOI:** 10.1101/2024.05.23.595639

**Authors:** Dilay Hazal Ayhan, Serena Abbondante, Domingo Martínez-Soto, Siyuan Wu, Ricardo Rodriguez-Vargas, Shira Milo, Katherine Rickelton, Vista Sohrab, Shunsuke Kotera, Tsutomu Arie, Michaela Ellen Marshall, Marina Campos Rocha, Sajeet Haridas, Igor V. Grigoriev, Neta Shlezinger, Eric Pearlman, Li-Jun Ma

## Abstract

*Fusarium oxysporum* is a cross-kingdom pathogen. While some strains cause disseminated fusariosis and blinding corneal infections in humans, others are responsible for devastating vascular wilt diseases in plants. To better understand the distinct adaptations of *F. oxysporum* to animal or plant hosts, we conducted a comparative phenotypic and genetic analysis of two strains: MRL8996 (isolated from a keratitis patient) and Fol4287 (isolated from a wilted tomato [*Solanum lycopersicum*]). Infection of mouse corneas and tomato plants revealed that, while both strains cause symptoms in both hosts, MRL8996 caused more severe corneal disease in mice, whereas Fol4287 induced more pronounced wilting symptoms in tomato plants. *In vitro* assays using abiotic stress treatments revealed that the human pathogen MRL8996 was better adapted to elevated temperatures, whereas the plant pathogen Fol4287 was more tolerant to osmotic and cell wall stresses. Both strains displayed broad resistance to antifungal treatment, with MRL8996 exhibiting the paradoxical effect of increased tolerance to higher concentrations of the antifungal caspofungin. We identified a set of accessory chromosomes (ACs) that encode genes with different functions and have distinct transposon profiles between MRL8996 and Fol4287. Interestingly, ACs from both genomes also encode proteins with shared functions, such as chromatin remodeling and post-translational protein modifications. Our phenotypic assays and comparative genomics analyses lay the foundation for future studies correlating genotype with phenotype and for developing targeted antifungals for agricultural and clinical uses.

**Importance:** *Fusarium oxysporum* is a cross-kingdom fungal pathogen that infects both plants and animals. In addition to causing many devastating wilt diseases, this group of organisms was recently recognized by the World Health Organization as a high-priority threat to human health. Climate change has increased the risk of *Fusarium* infections, as *Fusarium* strains are highly adaptable to changing environments. Deciphering fungal adaptation mechanisms is crucial to developing appropriate control strategies. We performed a comparative analysis of *Fusarium* strains using an animal (mouse) and plant (tomato) host and *in vitro* conditions that mimic abiotic stress. We also performed comparative genomics analyses to highlight the genetic differences between human and plant pathogens and correlate their phenotypic and genotypic variations. We uncovered important functional hubs shared by plant and human pathogens, such as chromatin modification, transcriptional regulation, and signal transduction, which could be used to identify novel antifungal targets.

## INTRODUCTION

*Fusarium oxysporum*, a cross-kingdom pathogen, is included in the list of health-threatening fungi by the World Health Organization (WHO) (1) and is considered to be among the top five most important plant pathogens (2). Corneal infections (keratitis) caused by *Fusarium* pathogens are an important cause of blindness worldwide, resulting in over one million new cases of blindness annually (3, 4). Indeed, *Fusarium* spp. are the most common cause of fungal keratitis in India (5, 6), China (7, 8), South Africa (9), and Brazil (10). *F. oxysporum* was also the cause of the 2005–2006 keratitis outbreak among contact lens wearers in the US and other temperate regions of the world (11, 12). As a plant pathogen, *F. oxysporum* causes devastating vascular wilt diseases in many economically important crops, including tomato (*Solanum lycopersicum*) (13), cotton (*Gossypium hirsutum*) (14, 15), and banana (*Musa* sp.) (16), and is responsible for billions of dollars in annual yield losses. Disease severity caused by this cross-kingdom pathogen is compounded by the lack of effective drugs, as only a limited number of antifungal agents are available to control eukaryotic fungal pathogens, and most *Fusarium* isolates are notorious for their broad resistance to many of these drugs (17–19).

Host-specific virulence is well recognized among plant pathogenic *F. oxysporum* isolates, as a single *F. oxysporum* species complex (FOSC) member typically infects only one or two plant species and is recognized as a *forma specialis* among plant pathogens. Within the FOSC, over 100 recognized *F. oxysporum formae speciales* infect more than 100 diverse plant hosts (20). We previously identified horizontally inherited accessory chromosomes (ACs) that lack genes associated with housekeeping functions but are enriched for genes related to fungal virulence as determinants of host-specific pathogenicity within the FOSC (21–24). Therefore, ACs (also referred to as pathogenicity chromosomes) encode several functionally validated virulence factors toward plant hosts, such as Secreted in Xylem (*SIX*) effectors (25), transcription factors (26), and kinases (27, 28). The importance of *F. oxysporum* ACs was demonstrated by the finding that a non-pathogenic *F. oxysporum* strain became virulent upon receiving ACs from a pathogenic strain (21, 29, 30).

No specific *forma specialis* has been reported for clinical isolates of *F. oxysporum*. The conventional wisdom was that disseminated fusariosis is the result of opportunistic infection of immunocompromised patients by environmental isolates, and corneal infections occur as a result of ocular trauma or contaminated contact lenses. The finding that a plant pathogenic isolate was able to infect immunosuppressed mice and cause systemic disease supports this notion (31). However, we reported that distinct sets of ACs were reported in the genomes of two clinical FOSC strains (32) and found to lack the genomic signatures that define plant pathogenic ACs, including *SIX* effector genes and some repeat elements commonly present upstream of these genes (33). By contrast, these two clinical isolates share AC regions that are enriched in genes encoding metal ion transporters and cation transporters (32).

Using the well-established murine model of fungal keratitis (4) and the tomato vascular wilt models (25), this study compared the human pathogenic strain MRL8996 isolated from a keratitis patient and the plant pathogenic strain Fol4287 isolated from a diseased tomato to test the hypothesis that distinct ACs contribute to the unique adaptation of fungal strains to animal or plant hosts. Both strains exhibited greater virulence in their respective hosts. Although we observed cross-kingdom virulence, these interactions resulted in milder disease in plants and in animals. These observations were supported by *in vitro* studies showing that MRL8996 was better adapted to higher temperatures, while Fol4287 tolerated conditions imposing osmotic or cell wall stress. These phenotypic assays revealed the unique adaptations of human and plant pathogens, providing a platform to dissect their different interactions with diverse hosts. Comparative genomics highlighted distinct genetic elements unique to each strain, offering potential testable hypotheses to correlate phenotypic to genotypic variations between a human and a plant pathogen. We also uncovered important functional hubs in ACs used by both human and plant pathogens, including chromatin modification, transcriptional regulation, and signal transduction, potentially identifying novel antifungal targets.

## RESULTS

### COMPARATIVE PHENOMICS REVEALS HOST-SPECIFIC ADAPTATION

We selected two *F. oxysporum* strains, one isolated from a keratitis patient and one from a tomato plant showing wilting symptoms as representative human and plant pathogens. MRL8996 was originally isolated from an infected cornea of a patient in the 2005–2006 contact lens–associated keratitis outbreak cohort (34). In addition, the genome of this strain has been published (35) and a clinically relevant mouse model of keratitis has been established (36). Phylogenetically, this keratitis strain is grouped with other human pathogenic isolates that cause systemic disease (37, 38). The tomato pathogenic strain Fol4287 was originally isolated from a diseased tomato plant in Spain (39, 40) has been adopted by the international Fusarium research community as a reference strain (21).

### MRL8996 is more virulent than Fol4287 in infected mouse corneas

To identify potential differences in virulence between Fol4287 and MRL8996 during corneal infection, we infected corneas of C57BL/6 mice with 5,000 *Fusarium* conidia from each strain (inoculum concentrations from 5,000 to 20,000 conidia are shown in Supplementary Fig. S1). After 24 h, we observed significantly higher corneal opacification MRL8996-inoculated eyes (**Fig. 1A, B**), compared to Fol428-inoculated corneas (**Fig. 1A, B**). We also examined fungal viability following eye infection with either fungal strain and detected significantly more colony-forming units (CFUs) in corneas infected with MRL8996 compared to Fol4287 (**Fig. 1C, Fig. S1**), indicating that the host was more efficient at containing the infection and possibly killing the Fol4287 strain compared to MRL8996.

**Figure 1.**
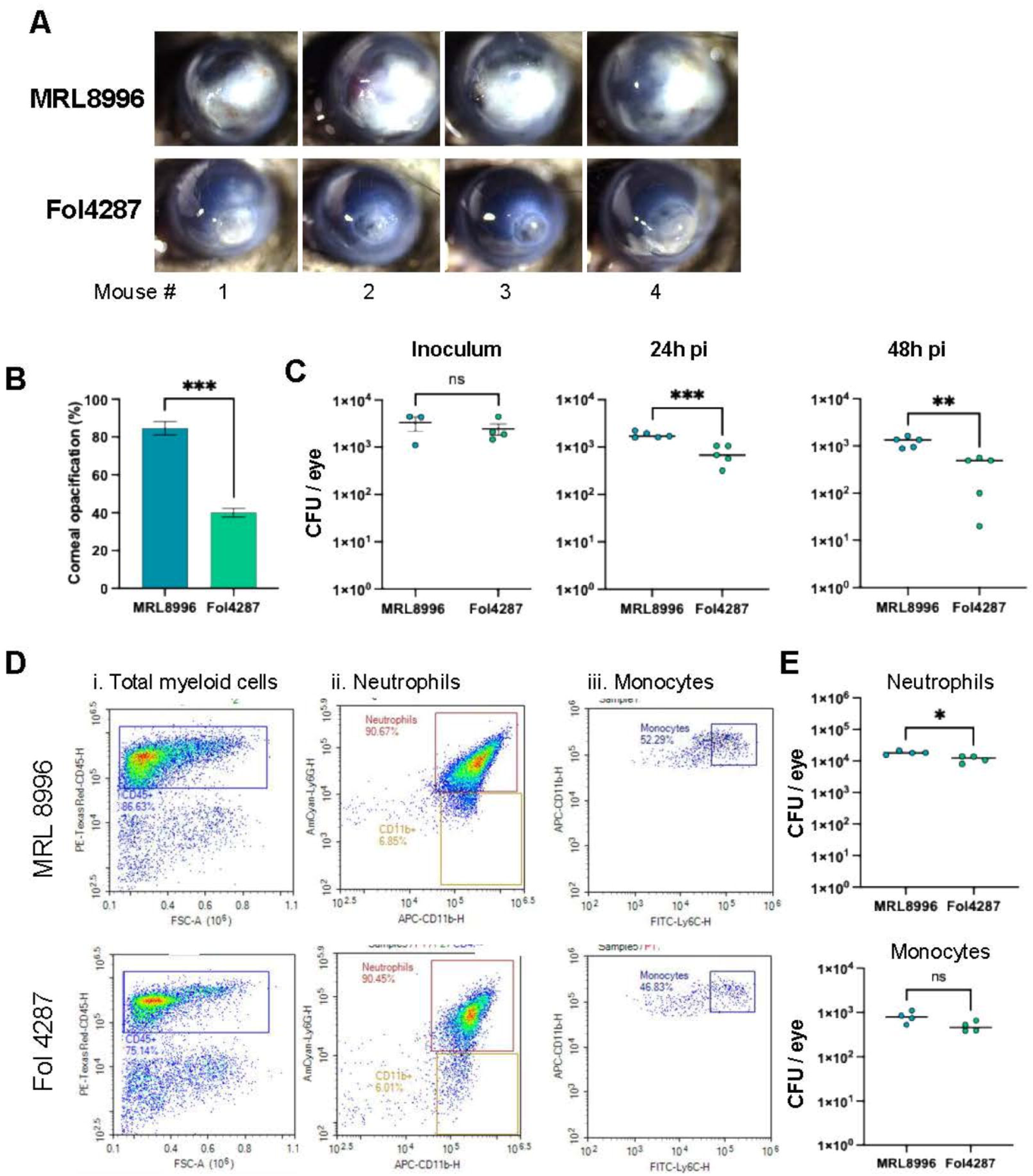
*In vivo* pathogenicity of clinical (MRL8996) and agricultural (Fol4287) *F. oxysporum* strains in infected corneas. **A.** Representative mouse corneas (n = 4) 24 h after intrastromal injection of 5,000 swollen conidia. **B.** Disease severity, as measured by the corneal opacification (94) (\*\**\* p* < 0.001). **C.** Viable conidia from infected corneas at t = 0 (inoculum; 5,000 conidia) and after 24 h, as indicated by the number of colony-forming units (CFUs) (*\*\* p* < 0.01). **D.** Infiltrating neutrophils and monocytes identified by flow cytometry. Total cells were isolated from collagenase-digested corneas at 24 h post-infection (hpi) with MRL8996 (top panels) or Fol4287 (bottom panels). Cells were identified using (i) CD45 for total myeloid cells (ii); CD45+ Ly6G+ and CD11b+ to identify neutrophils; and (iii) CD11b+ Ly6C+ monocytes. **E.** Quantification of neutrophils (top) and monocytes (bottom) in multiple infected corneas at 24 hpi. Significance was determined by a paired Student’s *t*-test, where * *p* < 0.05 was considered significant.

The major cause of corneal disease following fungal infection is the recruitment of CD45+ myeloid cells, including neutrophils and monocytes to the corneal stroma (4). To quantify these cells following infection with each *F. oxysporum* strain, we dissected corneas that were infected with MRL8996 or Fol4287, digested them with collagenase, and incubated the infiltrating cells with antibodies against CD45 and CD11b (which recognize total myeloid cells) and Ly6G and Ly6C (which identify neutrophils and monocytes, respectively) (41). Consistent with disease severity, corneas infected with MRL8996 had a higher percentage of CD45+ myeloid cells than those infected with Fol4287 (86.6% and 75.1%, respectively) (**Fig. 1D-i**). However, ∼90% of CD45+ myeloid cells were Ly6G+ neutrophils in corneas infected with each fungal strain (**Fig. 1D-ii**). Due to the higher percentage of CD45+ myeloid cells, we observed significantly more Ly6G+/CD11b+ neutrophils in corneas infected with the keratitis strain MRL8996 compared to the plant pathogenic strain Fol4287 (**Fig. 1E**). There was no significant difference in the number of Ly6C+/CD11b+ monocytes (**Fig. 1D-iii, E**).

Collectively, these findings indicate that the clinical isolate MRL8996 causes more severe disease and survives better than the plant pathogen Fol4287 in mouse corneas.

### Fol4287 is more virulent than MRL8996 in infected tomato plants

To examine the relative disease severity caused by these two *F. oxysporum* strains in plants, we inoculated tomato seedlings with MRL8996 or Fol4287 using a well-established wilting disease assay (42–44). Briefly, we soaked cleaned roots of tomato seedlings in a 10^6^ spores/mL suspension for 45 min before gently transplanting the inoculated seedlings in sterilized soil. As a mock control, roots were incubated in sterile distilled water. We used a disease index scale from 1 to 5 to measure disease severity (45), with 1 for healthy plants; 2 for plants showing wilted leaves with chlorotic areas; 3 for plants with necrotic spots; 4 for plants with wilted leaves and whole plants showing chlorosis, areas of necrosis, and defoliation; and 5 for dead plants.

At 10 days post-inoculation (dpi), the mock-inoculated seedlings were completely healthy, with a disease index of 1. By contrast, all tomato seedlings infected with the plant pathogen Fol4287 developed severe wilt symptoms, with an average disease index of 4, with wilting leaves showing areas of chlorosis, necrosis, and plant defoliation. Although the keratitis strain MRL8996 also caused chlorosis, the disease symptoms were significantly less pronounced, with an average disease index of 2 (**Fig. 2A, B**). The distinction between these strains was even more pronounced when tracking fungal colonization by confocal microscopy after wheat germ agglutinin (WGA)-Alexa Fluor 488 (green) staining of fungal cell walls and propidium iodide staining (red) of plant cell walls (46). As reported for wilt pathogens (44), the plant pathogenic strain Fol4287 invaded the xylem tissues in both primary and lateral roots at 4 dpi. By contrast, the keratitis strain MRL8996 was restricted to the epidermis and part of the cortex cell layers in both primary or lateral roots (**Fig. S2**). Furthermore, we documented that only Fol4287 colonized the vascular vessels of primary roots, lateral roots, and stems (**Fig. 2E**) and could be re-isolated from inoculated plant stems (**Fig. 2 C, D**).

**Figure 2.**
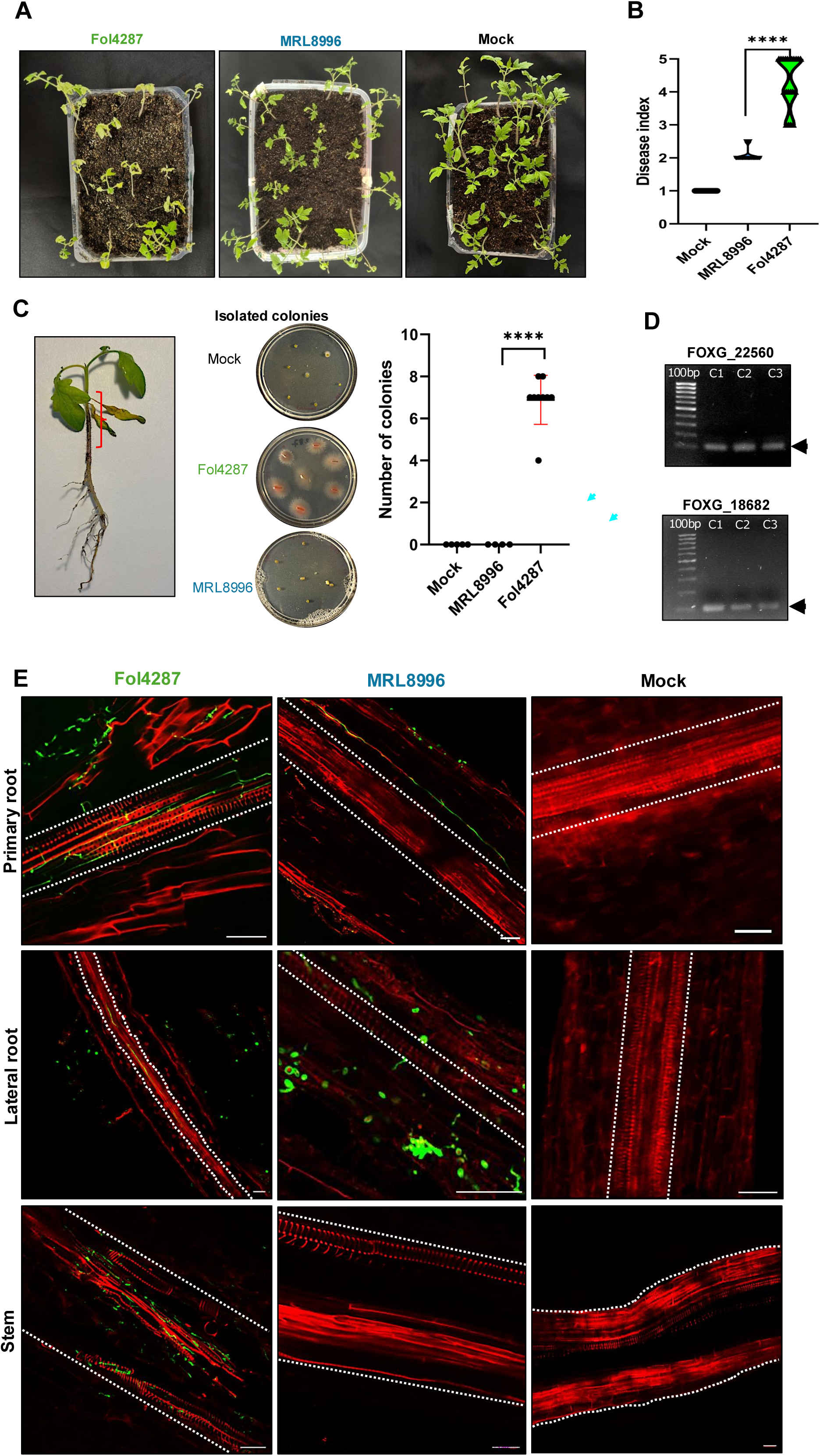
*In vivo* pathogenicity assay of clinical (MRL8996) and agricultural (Fol4287) *F. oxysporum* strains using the tomato vascular disease model. **A.** Representative photographs of tomato plants inoculated with Fol4287, MRL8996, or water (mock treatment). Ten-day-old tomato seedlings were infected and imaged 10 days later. **B.** A disease index of 1–5 was used to measure disease severity as described previously (45): 1 for healthy plants; 2 for plants showing wilted leaves with chlorotic areas; 3 for plants with necrotic spots; 4 for plants with wilted leaves and whole plants showing chlorosis, areas of necrosis, and defoliation; and 5 for dead plants. **C.** Re-isolation of fungal inoculum from inoculated plant stems. The red bracket indicates the area of the plant stem used for fungal isolation. Small slides of the stem were inoculated in Petri dishes with PDA medium. A total of 6 plants from each treatment were used for this assay. We isolated fungal colonies only from the stems of tomato plants inoculated with Fol4287. **D**. PCR amplification of 100 bp performed for two specific genes (FOXG_22560 and FOXG_18682) of Fol4287 in three different isolated colonies. **E**. Tracking the colonization of Fol4287, MRL8996, and mock infection using confocal microscopy. Fungal hyphae (in green) were stained with WGA-Alexa Fluor 488 and detected by excitation at 488 nm and emission at 500 to 540 nm. Plant tissues (in red) were stained with propidium iodide and detected with excitation at 561 nm and emission at 580–660 nm. Xylem is indicated by the dotted lines. The primary roots, the lateral roots, and the stems are visualized top to bottom.

Overall, these findings demonstrate that the plant pathogen caused significantly greater disease severity in tomato seedlings than the clinical pathogen.

### MRL8996 and Fol4287 exhibit resistance to different abiotic stress conditions

To explore cross-kingdom adaptation to different environmental conditions, we exposed the two *F. oxysporum* strains to the abiotic stresses of high salinity (0.6 M NaCl), oxidative stress (1 mM H_2_O_2_), and cell wall stress (1 mg/mL Congo Red), or to different temperatures (28°C or 34°C) and pH (5.0 or 7.4). The higher temperature and pH were chosen since they reflect the conditions of the human cornea (47). We calculated the growth rates of each strain at 3 dpi under each condition as the slope of the growth curve (diameter of the colony/dpi). Notably, the keratitis strain MRL8996 formed larger colonies than the plant pathogen Fol4287 in rich and minimal media at both temperatures tested whereas Fol4287 had a slower growth rate (**Fig. 3A, Table S1**, **Fig. S3A**).

**Figure 3.**
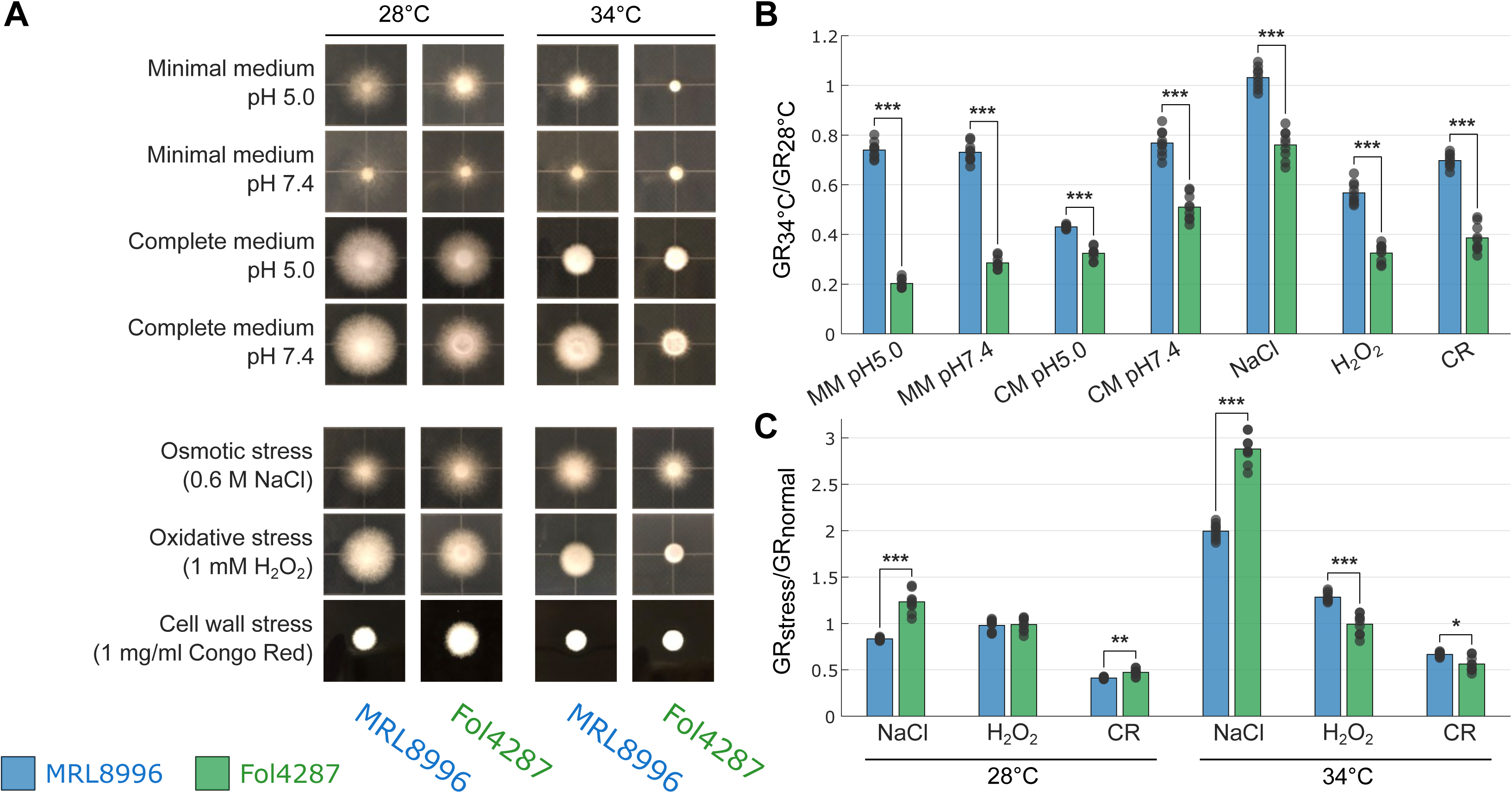
*In vitro* growth of clinical (MRL8996) and agricultural (Fol4287) *F. oxysporum* strains subjected to abiotic stress. **A.** Colony morphology of MRL8996 and Fol4287 on minimal medium (modified Czapek-Dox agar) pH ∼5, minimal medium pH 7.4, complete medium (CM, yeast extract peptone dextrose [YPD] agar), pH ∼5, complete medium pH 7.4, osmotic stress medium (potato dextrose agar [PDA] containing 0.6 M NaCl), oxidative stress medium (PDA with 1 mM hydrogen peroxide [H_2_O_2_]), and cell wall stress medium (YPD with 1 mg/mL Congo Red). The plates were incubated at 28°C or 34°C. The images are representative of three replicates and were taken at 2 days post-inoculation **B.** Ratio of the growth rates (GRs) of colonies under the same conditions as in **A**. at 34°C and 28°C (GR_34°C_/GR_28°C_) as a representation of temperature adaptation. **C.** Ratio of mean rates of growth under stress conditions (GR_stress_) and in CM (GR_normal_) as a representation of stress tolerance at 28°C and 34°C. Data points indicate each growth rate value at 34°C normalized to each growth rate value at 28°C, with each bar showing the mean value. * *p* < 0.05, ** *p* < 10^−5^; *** *p* < 10^−9^, calculated by a two-sample *t*-test.

To examine the tolerance of these strains to different stresses, we calculated the ratio of their growth rates under two different conditions. For instance, to assess tolerance to different temperatures, we calculated the ratio of the fungal growth rates between 34°C and 28°C (GR_34°C_/GR_28°C_). While both strains grew more slowly at the higher temperature, the growth rate of the plant pathogen Fol4287 was significantly slower at this temperature compared to the keratitis strain MRL8996 under all conditions tested **(Fig. 3B, C)**. At 34°C, lowering the pH from the human physiological pH (7.4) to an acidic pH (5) environment decreased the growth of the keratitis strain by 40%. Our observation that the human pathogenic strain MRL8996 has better tolerance to an elevated temperature of 34°C at pH 7.4 may reflect the adaptation of MRL8996 to the mammalian environment.

Conversely, the plant pathogenic strain Fol4287 exhibited significantly higher tolerance to osmotic stress (0.6 M NaCl) than MRL8996 (**Fig. 3C, Fig. S2B**). Compared to growth in complete medium alone at 28°C, the growth of MRL8996 in the presence of 0.6 M NaCl decreased by 16.7% relative to the control condition, whereas the growth of strain Fol4287 increased by 22.7%, revealing a significant difference in their response to osmotic stress (*p* < 10^-5^). At 34°C, both strains grew better under higher salinity conditions compared to the complete medium alone. While growth of MRL8996 increased by 99.4% under these conditions, growth of Fol4287 increased by 187.6% (*p* < 10^-9^). Therefore, the plant pathogenic strain Fol4287 is better adapted to higher salinity conditions than MRL8996 at both temperatures (**Fig. 3C, Fig. S2B**).

Surprisingly, 1 mM H_2_O_2_ treatment (to induce oxidative stress) did not affect the growth rates of either strain at 28°C. However, the human pathogenic strain MRL8996 exhibited resistance to oxidative stress, with a 28.4% increase in growth at 34°C compared to samples grown in complete medium alone. Further, cell wall stress imposed by the addition of 1 mg/mL Congo Red to the medium inhibited the growth of both strains. At 28°C, compared to growth in YPD medium alone, the growth rates of MRL8996 and Fol4287 were significantly lower (by 58.8% and 52.9%, respectively). At 34°C, the growth rate was reduced by 33.3% for MRL8996 and 43.8% for Fol4287 compared to the control.

We repeated *in vitro* phenotyping of the two reported strains and added one additional plant pathogenic isolate Fo5176 (infecting *Arabidopsis thaliana*), and one additional clinical isolate NRRL 32931 (isolated from a leukemia patient) (**Fig. S3, Table S1**) and observed consistent results. While growth rates were reduced for all four strains at 34°C compared to 28°C, the reduction of the two plant strains was significantly greater than the two clinical isolates. MRL8996 exhibited the highest tolerance to heat compared with Fo5176. The clinical isolate NRRL 32931 grew significantly better at physiological pH (7.4) compared to the acidic pH (5) for both Minimal Media and Complete Media (**Fig. S3B**). Also consistent with the previous findings, the plant isolates show higher resistance to high salinity in both temperatures compared to the two human clinical isolates. Fo5176 surprisingly shows high resistance to salinity and cell wall stress under both temperatures, with the difference more pronounced at 34°C (**Fig. S3C**).

Together, these findings indicate that the clinical isolate MRL8996 is better adapted to the physical conditions of the animal host, such as elevated temperature, whereas the plant pathogenic strain Fol4287 is more tolerant to increased salinity. These findings reflect a complex, interconnected regulatory relationship between the physiological adaptation of the fungus and fungal–host interactions.

### MRL8998, but not Fol4287, exhibits tolerance to high caspofungin concentrations

Caspofungin acetate (CFA) is an antifungal agent which targets 1,3-β-glucan synthase that mediates biosynthesis of one of the main components of the fungal wall (48, 49). However, different fungi show diverse responses to CFA, as clinical isolates of *Candida albicans* and *Aspergillus fumigatus* are sensitive to CFA (50), whereas some less common fungal human pathogens, including *Fusarium* spp., are resistant to clinically relevant levels of CFA (51).

In the current study, the keratitis strain MRL8998 exhibited enhanced tolerance to high caspofungin concentrations compared with Fol4287 (**Fig. 4A, Fig. S4**). The growth rate of the plant pathogen Fol4287 decreased from 80% to 70% with an increase in caspofungin concentration from 0.2 µg/mL to 8 µg/mL compared to growth in the absence of caspofungin, showing a strong tolerance to caspofungin without a clear dose-dependent response. In contrast, the growth rate of the keratitis strain MRL8996 dropped by 50% under 2 µg/mL caspofungin, with a further reduction to a 35% decrease at 8 µg/mL, relative to the control (**Fig. 4B**). This paradoxical caspofungin effect was also observed when measuring the time required to reach 50% conidial germination (**Fig. 4C**). While there was no significant difference in the time needed for the plant pathogen Fol4287 to reach a 50% germination rate in the presence of caspofungin, the human keratitis strain required a significantly longer time to reach 50% gemination under 0.5 µg/mL caspofungin treatment compared to the no caspofungin control or 8 µg/mL caspofungin treatment. Together, these findings indicate the different responses toward the antifungal caspofungin.

**Figure 4.**
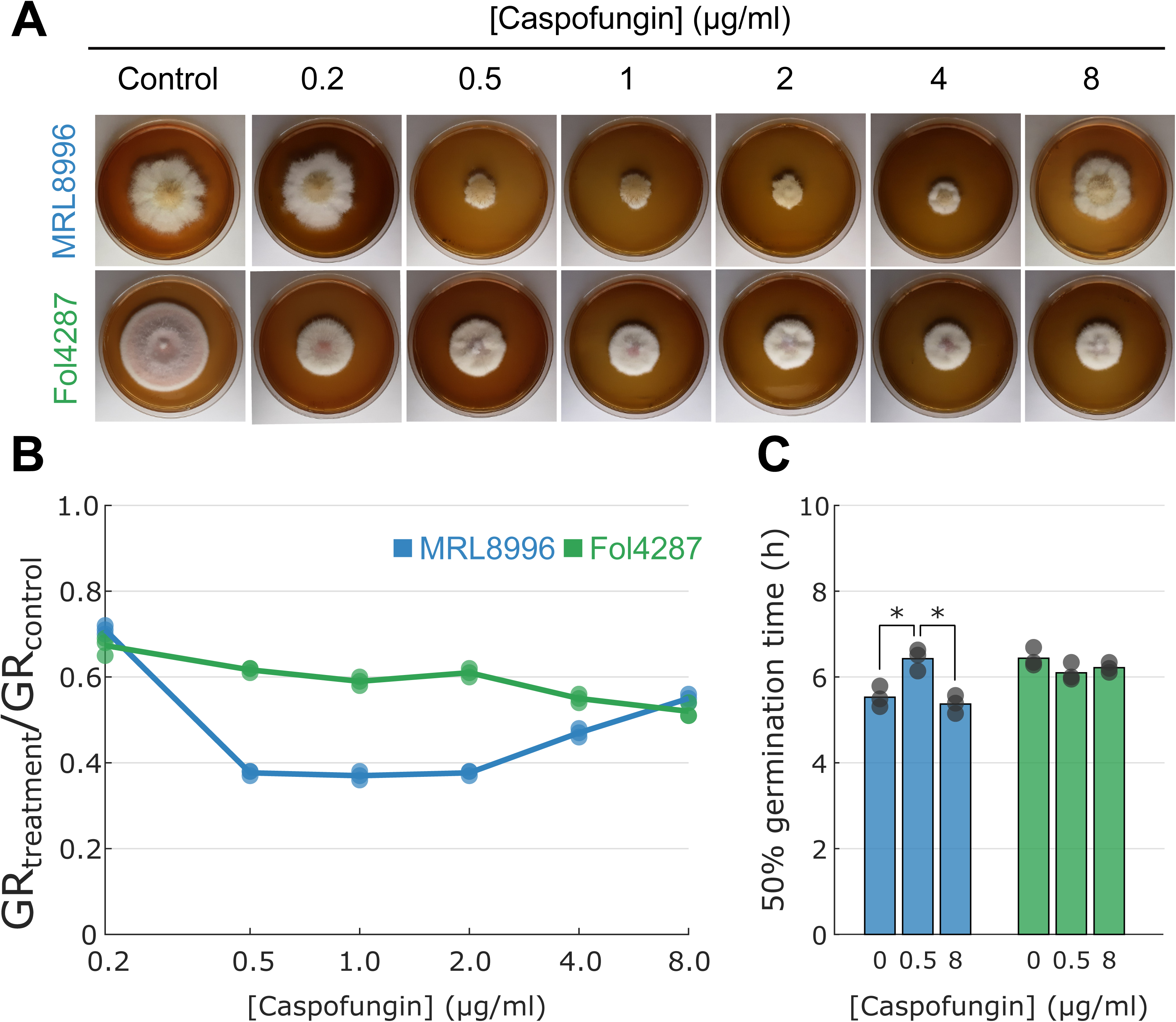
*F. oxysporum* keratitis strain MRL8896 exhibits a paradoxical effect to caspofungin. **A.** Representative images of radial fungal growth in the presence of caspofungin. **B.** Ratio of growth rates (GRs) for colonies in the presence of caspofungin (treatment) and in the absence of caspofungin (control). **C.** Effect of caspofungin concentration on germination time. For each strain, 1 × 10^4^ conidia were inoculated onto glass coverslips containing 200 µL of liquid PDA medium and incubated at 28°C for 24 h. Two hundred conidia were counted, and the percent germination was calculated at 0, 2, 4, 6, 8, and 12 h. The time when 50% of the spores germinated was calculated based on the closest datapoint in a log-logistic regression.

### COMPARATIVE GENOMICS REVEALS STRAIN-SPECIFIC ACCESSORY CHROMOSOMES WITH DISTINCT TRANSPOSON PROFILES

We used a contour-clamped homogeneous electric field (CHEF) gel to observe the diversity of karyotypes among *F. oxysporum* genomes and confirmed the presence of ACs in Fol4287 (21, 52), MRL8996 and NRRL 32931 (32) (**Fig. 5A**). Comparative genomics confirmed the conservation of core chromosomes (CCs) (**Fig. 5B**, **Table S2**) and revealed three small CCs (chromosomes 11, 12, and 13) that were less conserved, with an average of 94.0% sequence identity over 35.1% coverage. These findings support the three-speed evolutionary hypothesis (53). The conservation of CCs enabled us to identify ACs and sequences unique to the plant and human strains. As highlighted in darker blue and darker green boxes in **Fig. 5B**, the genomes of MRL8996 and Fol4287 contained 6.4 Mb and 9.8 Mb of accessory sequences corresponding to 12.8% and 18.7% of the total genome, respectively.

**Figure 5.**
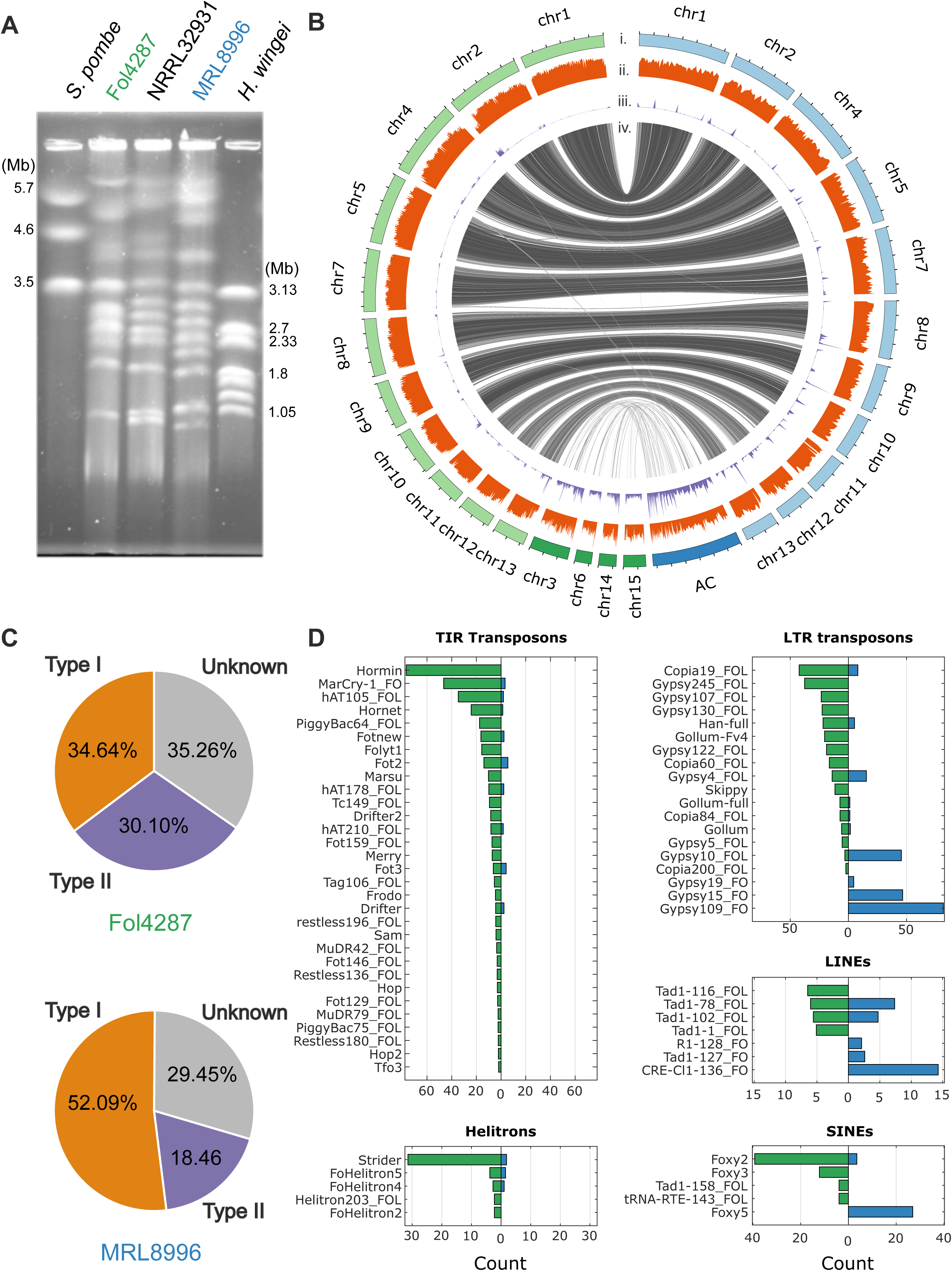
Compartmentalized genomes of the clinical (MRL8996) and agricultural (Fol4287) *F. oxysporum* strains. **A.** Contour-clamped homogeneous electric field (CHEF) gel of small accessory chromosomes (ACs) of the agricultural isolate Fol4287, the control clinical isolate NRRL32931, and the clinical isolate MRL8996. *Schizosaccharomyces pombe* (left) and *Hansenula wingei* (right) were used as markers. The band sizes are shown in Mb. ACs are typically below 2 Mb in size, except for two known ACs, chr3 and chr6 in Fol4287, which are much larger due to recent segmental duplications (21). **B.** Whole-genome comparison between MRL8996 and Fol4287 revealed 11 homologous core chromosomes (light green and light blue, respectively) and accessory sequences (dark green and dark blue, respectively) (i). The accessory sequences typically displayed low gene density (ii) and high repetitive sequence composition (iii). The syntenic alignment between MRL8996 and Fol4287 using nucmer (iv) indicates the accessory regions lacking synteny between MRL8996 and Fol4287. **C.** Distribution of all identified transposable I retrotransposons, class II DNA transposons, and unknown transposable elements. **D.** Transposon abundance in the Fol4287 and MRL8996 genomes. Class I retrotransposons include LTRs (long terminal repeats), LINEs (long interspersed nuclear elements), and SINEs (short interspersed nuclear elements). Class II DNA transposons are classified into TIRs (terminal inverted repeats) and Helitrons.

Our comparative analysis also revealed overall conservation of the mitochondrial genome (99.0%) and highlighted a divergent sequence around open reading frame 2285 (ORF2285) (**Fig. S3**), a known variable mitochondrial region in the genus *Fusarium* (54).

The repeat contents in the Fol4287 and MRL8996 genomes were similar, with 6.6% and 5.4% values, respectively (**File S1, S2**). Transposable elements (TEs), a signature of accessory sequences (55), were enriched in the ACs of both genomes (27.2% and 23.8%, respectively) (**Table S3**), but with distinct transposon profiles. In the Fol4287 genome, approximately one-third of identified TEs were type I retrotransposons, and the another third was type II DNA transposons. In the MRL8896 genome, more than half were identified as type I and less than 20% as type II TEs (**Fig. 5C**).

Some class I retrotransposons are conserved within the genus of *Fusarium* (21). Similarly, the long terminal repeat (LTR) transposon *Copia*, was present in both the Fol4287 and MRL8996 genomes (**Fig. 5D, Table S4**). However, some LTR elements with the highest copy numbers in MRL8996, such as several *Gypsy* elements, were not present in Fol4287. Similarly, the most abundant long interspersed nuclear element (LINE) transposon in MRL8996, the *Cnl1* CRE-type non-LTR retrotransposon, was not present in Fol4287. Of the three short interspersed nuclear element (SINE) transposon families, the *Foxy2* family was the most abundant in Fol4287, while the *Foxy5* family was the most abundant in MRL8996. Conversely, Fol4287 contained unique class II DNA transposons (**Fig. 5D, Table S4**).

The most abundant DNA TE in Fol4287, *Hormin* (a miniature *Hornet* element), was absent from the MRL8996 genome. Miniature impala elements (MIMPs) (33) were not present in MRL8996. Similarly, Fol4287 contained significantly more Helitrons (56) compared to MRL8996. There were also many unclassified repeat elements that were not shared between the Fol4287 and MRL8996 genomes (**Fig. S4**). The distinction between different TE profiles is important, as class I and class II TEs play different roles in the genome dynamics of MRL8996 and Fol4287.

### STRAIN-SPECIFIC ACCESSORY CHROMOSOMES CONTRIBUTE TO DISTINCT FUNCTIONAL ADAPTATIONS

We annotated the genome of MRL8996 using the Joint Genome Institute annotation pipeline (57) and assigned Gene Ontology (GO) terms to genes in both genomes using the Mycocosm GO annotation pipeline (**Table 1**). Among the 16,631 and 20,925 predicted protein-coding genes in the MRL8996 and Fol4287 genomes, respectively, we identified 2,017 (12.1%) and 3,890 (18.6%) genes located on their respective ACs. GO functional enrichments (*p-*value < 10^−3^) revealed some functional categories shared by both pathogens (**Fig. 6** category III), in addition to strain-specific functions (**Fig. 6** categories I and II, **Table S5, S6**).

**Figure 6.**
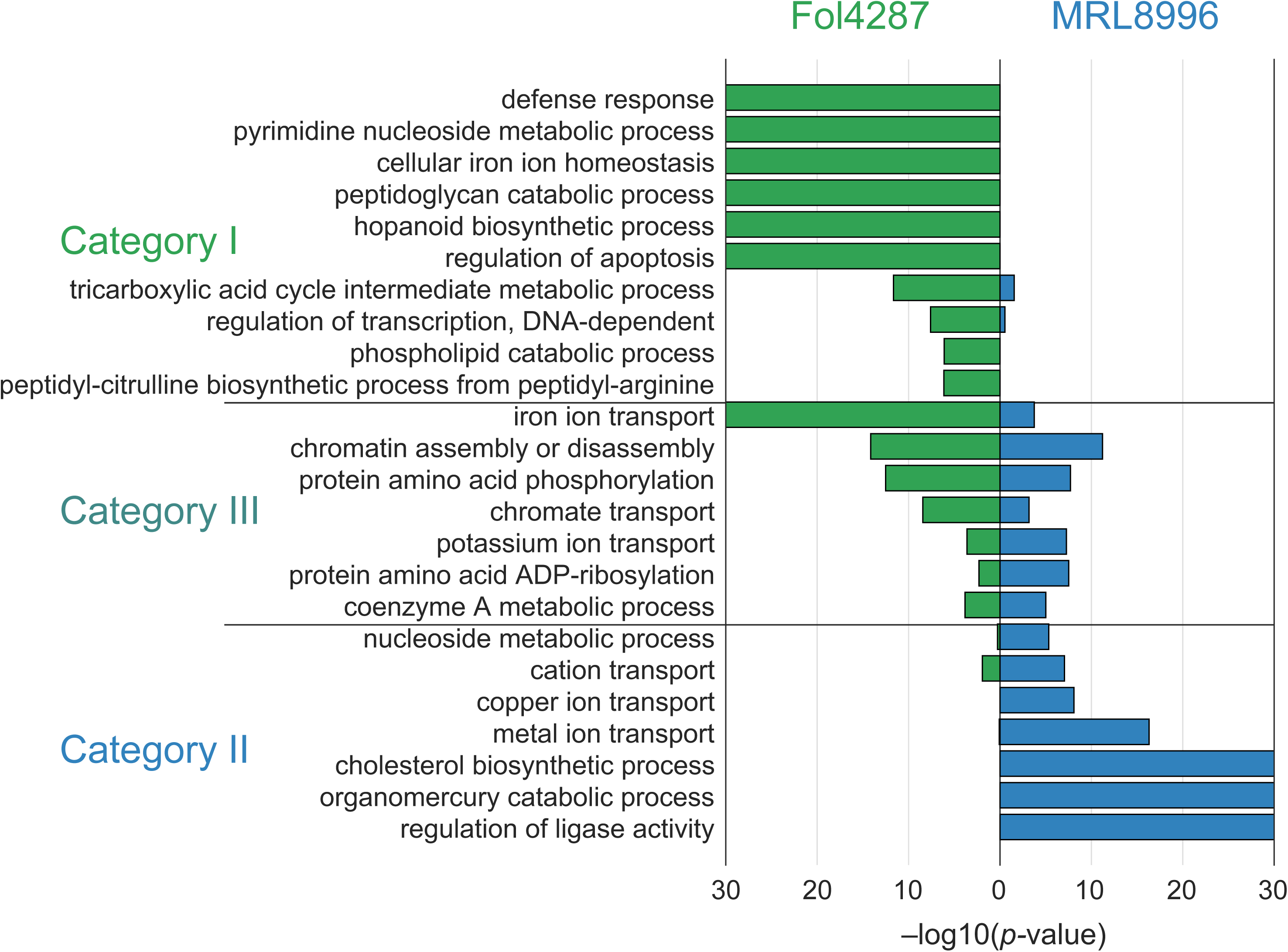
Enriched Gene Ontology terms among accessory genes of the clinical (MRL8996) and agricultural (Fol4287) *F. oxysporum* isolates. Enriched Gene Ontology (GO) terms harbored by the ACs from Fol4287 (green) and MRL8996 (blue). Category I represents GO terms enriched in Fol4287 ACs. Category II represents GO terms enriched in the ACs of both genomes. Category III represents GO terms enriched in MRL8996 ACs. For graphical representation, a *p*-value = 0 is shown as −log10(*p*-value) = 30.

**Table 1.** Comparative annotation of the Fol4287 and MRL8996 genomes.

|  | **Fol4287** | **MRL8996** |
| --- | --- | --- |
| Total genes | 20,925 | 16,631 |
| Genes with GO annotation | 9,783 | 7,958 |
| Total AC genes | 3,890 | 2,017 |
| AC genes with GO annotation | 1,788 | 542 |

### Functional groups enriched in both genomes due to different sets of AC genes

1. *Chromatin assembly or disassembly*. The most significantly enriched functional term in both genomes was chromatin assembly or disassembly (GO:0006333), with *p =* 6 × 10^−12^ for MRL8996 AC genes and *p =* 7 × 10^−15^ for Fol4287 AC genes. Most genes under this annotation encode CHROMO domain-containing proteins that bind to the H3K9 di/trimethyl modification on histone H3, a hallmark of heterochromatic regions (58, 59) that are typically involved in the assembly or remodeling of chromatin and associated proteins or RNAs (60). Phylogenetic analysis grouped these genes into three clades (**Fig. S5**). The first clade shares homologs of CHROMO domain-containing protein 2 (Chp2) from fission yeast (*Schizosaccharomyces pombe*). The second clade encodes homologs of putative Heterochromatin protein 1 (HP1) (A0A0C4BKY0) based on AlphaFold prediction (61). The third clade includes genes encoding Zinc finger C2H2-type domain-containing proteins. Notably, Chp2 and HP1 both interact with chromatin in DNA–protein complexes (62) and functioning as a key regulator coordinating eukaryotic chromatin compaction, HP1 can bind to heterochromatin marks and recruits other factors to promote heterochromatin formation (63). *F. oxysporum* ACs are primarily composed of facultative heterochromatin (64). The equal expansion in this protein family suggests that establishing the ability to effectively open or close AC regions, either to exploit transcriptional networks or to allow genome maintenance activities, likely plays a crucial role in the functional regulation of *F. oxysporum* ACs.
2. *Post-translational modifications.* The other significantly enriched term (*p=* 1.95 × 10^−8^ for MRL8996 AC genes and *p =* 3 × 10^−13^ for Fol4287 AC genes) was protein phosphorylation (GO:0006468), an important cellular regulatory mechanism through kinases. This result was expected, as we previously identified a positive correlation between the total number of protein kinases encoded in a genome and the size of the proteome of an organism (65). However, each genome has its own distinct set of encoded proteins. For instance, the major AC-enriched kinases in MRL8996 are Halotolerance protein 4-like (HAL4) serine/threonine kinases (which regulate ion transporters), Calmodulin Dependent Protein Kinase Kinase 2 (which is involved in energy balance and perhaps mobile DNA repair activity), and Cell Division Cycle (CDC)-like kinases (which function in chromatin remodeling and DNA metabolism/repair). The kinases encoded by the AC of Fol4287 include the second copy of a TOR kinase paralog (a top regulator that dictates cellular stress responses) (65); choline kinase (a conserved APH superfamily member involved in antibiotic resistance); a TOMM system kinase/cyclase fusion protein; and the serine/threonine kinase protein kinase B (involved in stress responses and signal transduction).

In addition to phosphorylation, the other shared GO term involved in the post-translational modification of proteins is protein ADP-ribosylation (GO:0006471) (*p =* 2.99 × 10^−8^ for MRL8996 AC genes and *p =* 5 × 10^−3^ for Fol4287 AC genes). Several transporter genes, including a chromate transporter gene (GO:0015703), genes enriched in the terms potassium ion transport and iron ion transport, and a hydroxymethylglutaryl-CoA reductase gene (GO:0015936), the rate-limiting enzyme for ergosterol biosynthesis, were also enriched in both genomes (**Fig. 5C**, **Table S5**, **S6**).

### Fol4287 AC genes are uniquely enriched for defense responses and signaling

Among the best characterized Fol4287 AC genes are *SIX* effector genes, encoding *bona fide* fungal virulence factors (66–68) that are directly involved in fungal–plant interactions. Many genes identified in this study were reported previously. In addition, this study revealed a significant enrichment of Fol4287 AC genes in the GO term defense response (GO:0006952) compared to the core genes and the AC genes from MRL8896 **(Figure 6** category I, **Table S6)**. The AC genes in this group include two chitinase II genes and one patatin-like phospholipase gene involved in the manipulation of plant resistance proteins (69).

### MRL8996 AC genes are uniquely enriched for metal ion transport and catabolism

Whereas MRL8996 ACs lack *SIX* effector genes and genes encoding plant cell wall–degrading enzymes, unlike plant pathogenic *F. oxysporum* strains (32), other effectors are also detected. Also, MRL8996 ACs are uniquely enriched in genes involved in the transport and catabolism of metal ions such as copper, zinc, magnesium (GO:0006825 and GO:0030001), and mercury (GO:0046413) **(Figure 6** category II**, Table S5)**. Nine genes are specifically involved in copper ion transport, with seven Ctr-type copper transporter paralogs and two copper-exporting P-type ATPase (CopA) paralogs encoding proteins containing multiple copies of the copper chaperone (CopZ) domain. Other genes included in these categories encode a CorA-like magnesium/zinc transporter, with multiple copies of ankyrin repeats, and a Zrt-like, Irt1-like protein (ZIP)-type zinc ion transporter. Most of these genes are present in the genome of another clinical isolate, NRRL32931 (25 out of 33 genes). One of these shared genes encodes a mammalian ceruloplasmin homolog with two multicopper oxidase domains. Homologous sequences were previously detected among a few other opportunistic fungal pathogens and bacteria isolated from extreme environments, but not in plant pathogenic *Fusarium* species (32), suggesting that a host-specific gene network evolved in the human pathogens. Another GO term that was exclusively enriched in the MRL8996 ACs is regulation of ligase activity (GO:0051340).

Taken together, our results provide strong evidence that different ACs acquired by a plant or a human pathogen perform shared functions that are essential for chromatin modifications, transcriptional regulation, and post-translational modification of proteins. These essential regulatory mechanisms, which are involved in environmental sensing and cellular signal transduction, may hold key to understanding the crosstalk between the core and AC genomic regions. However, the strain-specific AC gene repertoires are uniquely enriched for defense responses and signaling in the plant pathogen and metal ion transport and catabolism in the human pathogen. We propose that these unique adaptations are important for fungal survival in different environments and hosts.

## DISCUSSION

The FOSC, a group of cross-kingdom fungal pathogens, includes both plant and animal pathogens. Our *in vivo* assays using animal and plant hosts confirmed the host specificity of the keratitis strain MRL8996 and the tomato pathogen Fol4287, as corneal infection with the keratitis strain resulted in significantly more corneal ulceration, while the plant pathogen caused significantly more wilt symptoms among all inoculated tomato seedlings. Consistent with host-specific adaptations, the keratitis strain MRL8996 grew significantly better at elevated temperature, whereas the tomato pathogen Fol4287 exhibited more tolerance to osmotic and cell wall stress. As a line of fungal adaptation, a human pathogen must overcome the host defenses associated with mammalian endothermy and homeothermy, while a plant pathogen must handle different stresses (70, 71). Our findings reveal interconnected fungal responses toward biotic stress from plant and animal hosts and abiotic stress in distinct environments. The distinct phenotypes observed in our *in vivo* and *in vitro* assays, together with the identification of unique ACs in each genome, lay the foundation for dissecting cross-kingdom fungal pathogenicity using this comparative model.

The widespread occurrence of the caspofungin paradoxical effect underscores the complexity of antifungal drug responses in filamentous fungi and provides an early warning about the potential complexity in treating infectious diseases caused by *Fusarium* species (72). In the current study, we detected the caspofungin paradoxical effect, but only for the keratitis strain MRL8996. Mechanisms underlying this phenomenon likely involve specific drug–target interactions for each strain(73), fungal stress responses (74), and cellular signaling pathways (72, 75–79). Future studies will identify underlying mechanisms that may lead to strategies to overcome the paradoxical effect and for improving the efficacy of antifungal therapies.

In addition, we observed severe disease phenotypes from *Fusarium* keratitis compared to other fungal infections. For instance, corneal infection with *Aspergillus fumigatus* requires 40,000 conidia as the inoculum, which still causes severe corneal disease (80). When we used the same titer of *F. oxysporum* conidia, almost all animals developed corneal ulceration and perforation within 24 h. Therefore, we lowered the inoculum to 5,000 conidia. This hypervirulence may contribute to *Fusarium* species being the leading cause of blindness among fungal keratitis patients (3, 5).

A unique advantage of our comparative system is the compartmentalized genomic structures for both pathogens, allowing us to characterize two distinct sets of ACs to be which revealed distinct transposon profiles: while DNA transposons dominate Fol4287 ACs, retrotransposons, especially LINE transposons, are highly abundant among MRL8996 ACs. Intriguingly, ACs from both genomes also encode proteins with shared functions, including chromatin assembly/disassembly, protein phosphorylation, and transcriptional regulation, which support our previous findings (65, 26) Our observation that there is shared enrichment of genes involved in chromatin assembly and disassembly points to the importance of chromatin remodeling regardless of host-specific functions. Identifying master regulators of the crosstalk between ACs and core components, especially those regulating the activities of both plant and animal pathogens, should uncover new targets for control of this cross-kingdom pathogen.

In summary, this is the first study documenting similarities and differences in the genotypes and phenotypes of two closely related pathogens: one infecting humans and one infecting plants. Our *in vivo* and *in vitro* assays allowed us to examine strain adaptation under differenet environmental stress conditions feasible. In addition to host-specific virulence, we also observed cross-kingdom virulence, as the human pathogen also colonized plant roots and the plant pathogenic strain infected corneas, although both resulted in milder disease symptoms. More in-depth research is needed on the molecular mechanisms underlying species divergence and adaptation among this group of pathogens.

## MATERIALS AND METHODS

### Fungal strains

The genome of *F. oxysporum* Fol4287 was first sequenced in 2010 (21), and an improved genome assembly was subsequently produced in 2018 (52). The strain is deposited in multiple public strain repositories, including the Fungal Genetics Stock Center (FGSC 9935), NCAUR/USDA (NRRL 34936), and CBS-KNAW (CBS 123668). The keratitis strain *F. oxysporum* MRL8996 was originally isolated in 2006 from the cornea of a patient with the contact lens–associated multistate outbreak fungal keratitis at Cleveland Clinic Foundation in Ohio, USA (34). The strain is available at NCAUR/USDA (NRRL 47514). MRL8996 is grouped with other human pathogenic isolates belonging to clade FOSC 3-a. The genome assembly for MRL8996 was produced using the same sequencing technologies and computational strategies used to assemble the Fol4287 genome to facilitate effective comparative analysis (32).

### Fungal growth conditions

The conidia of the fungal strains were stored at −80°C in an ultra-freezer in 25–50% (v/v) glycerol for long-term storage and propagated from stocks in liquid potato dextrose broth (PDB, BD Difco™, USA) or potato sucrose broth (PSB: 25% [w/v] boiled potato and 0.5% [w/v] sucrose) for at least 4 days in a shaking incubator at 28°C. The conidia were collected by filtering the liquid fungal cultures through two layers of Miracloth (EMD Millipore). The filtrate was centrifuged at 3700 g for 5 minutes and the spores were resuspended in sterile water to the desired concentration. The spore titers were determined using a hemocytometer.

### Mouse corneal infection

Fungal conidia were harvested from Sabouraud Dextrose Agar (SDA) medium and resuspended in phosphate buffered saline (PBS, pH 7.0). Mice were anesthetized by intraperitoneal injection with ketamine/xylazine. A 30-gauge needle was used to make a pocket in the corneal stroma, after which a 33-gauge Hamilton syringe was inserted, and 5 × 10^3^ conidia in 2 µl PBS was injected into the corneal stroma. Corneal opacity was photographed using a high-resolution stereo fluorescence MZFLIII microscope (Leica Micro-systems) and a Spot RT Slider KE camera (Diagnostics Instruments). All images were captured using SpotCam software (RT Slider KE; Diagnostics Instruments). Corneal opacity was quantified using ImageJ software. The experiments used 5 mice per group and were repeated 4 times.

### Colony-forming units (CFUs) from *Fusarium*-infected corneas

Infected whole eyes were homogenized in 1 mL of sterile PBS using a Retsch Mixer Mill MM300 (Qiagen, Valencia, CA) at 33 Hz for 4 min. Ten-fold dilutions were made, and the samples were plated on SDA plates and incubated at 34°C for 48 h. CFUs were determined by direct counting.

### Flow cytometry

Infected corneas were dissected, the vascularized iris was removed, and the corneas were incubated in 500 μL collagenase type I (Sigma-Aldrich) at a titer of 82 U per cornea for 1–2 h at 37°C. Cells were resuspended in 200 μL FACS buffer containing 4 μg/cornea Fc blocking Ab (anti-mouse CD16/32; eBioscience) on ice for 10 min, followed by incubation with anti-mouse antibodies against CD45, Ly6G CD11b, or Ly6C, all from BioLegend. Total cells were quantified using an ACEA Novocyte™ flow cytometer.

### Phenotyping of isolates

Yeast extract peptone dextrose (YPD) plates were used as complete medium, and modified Czapek-Dox plates were used as minimal medium. The pH levels of both media were adjusted to pH 7.4 using 6.5% (w/v) 0.1 M citric acid and 43.5% (w/v) 0.2 M Na_2_HPO_4_. YPD plates containing 0.6 M NaCl, 1 mM H_2_O_2_, and 1 mg/mL Congo Red were prepared for osmotic, oxidative, and cell wall stress treatments, respectively. Each isolate (Fol4387, MRL8996, NRRL 32931, and Fo5176) was grown in PSB for 4 days at 28°C, and the conidia were filtered as described above. Plates containing the same medium were inoculated with 2 μL of spore suspensions at concentrations of 5 × 10^6^, 5 × 10^5^, or 5 × 10^4^ conidia/mL, each in three replicates. The plates were incubated at 28°C or 34°C for 3 days and photographed once a day. The experiments were repeated twice.

### Statistical analysis of growth rates

The growth rates of each replicate and dilution were calculated as the slope of the 3-day growth curve. Five-way ANOVA with a linear model was performed for the following groups: strain, temperature, medium, dilution, and replicate, followed by a multi-comparison with strains, temperatures, and media groups to identify significant differences using MATLAB. To quantify adaptation to different temperatures, the growth rates of cultures with an initial inoculum of 1 × 10^3^ conidia were measured in triplicate. The values of the three replicates incubated at 34°C were normalized to the values of the three replicates incubated at 28°C (total of nine values). A two-way Student’s *t*-test was performed between Fol4287 and MRL8996 values. Similarly, to quantify the effects of stress treatment, the values of the three replicates of samples in medium containing different stress components were normalized to the values of the three replicates in control YPD medium (total of nine values), followed by a two-way Student’s *t*-test between Fol4287 and MRL8996 in MATLAB.

### Caspofungin sensitivity assay

Both strains were maintained in potato dextrose agar (PDA) medium. For solid medium, 2% (w/v) agar was added. The growth rate was determined by spotting 1 × 10^4^ conidia in the center of a 90-mm Petri plate containing 20 mL of solid PDA medium and incubating the plates at 28°C. The diameter was scored at 24-h intervals until 5 days (96 h) of incubation with three biological replicates. *To assess the strains’ germination kinetics*, 1 × 10^4^ conidia of each strain were inoculated onto glass coverslips containing 200 µL liquid PDA medium, and the samples were incubated at 28°C. A conidiospore was counted as germinated if it possessed a germ tube, which was readily detectable as a small protuberance on the spherical spore surface. Two hundred conidia were counted in each experiment. Times when 50% of conidia reach the stage of germination are recorded.

### Plant infection assay

Seeds of tomato (*Solanum lycopersicum*) cv. M82 were maintained in the dark at 4°C for 3 days. The seeds were surface sterilized in 70% (v/v) ethanol for 5 min and washed with 2.7% (w/v) sodium hypochlorite (NaClO) for 5 min. After removing the NaClO solution, the seeds were rinsed with sterile distilled water (SDW) three times. The seeds were then sown in pots filled with autoclaved soil (Promix BX) and watered with SDW. The soil was gently removed from the roots of 10-day-old tomato seedlings by rinsing them with abundant distilled water and SDW while avoiding root tissue damage. The roots of the seedlings were inoculated by dipping them into the respective human or plant pathogenic *Fusarium* spore suspension (10^6^ spores/ml) or in SDW as a mock infection control for 45 min before replanting the infected seedlings in soil. After infection, all seedlings were maintained in a growth chamber at 28°C under a 14-h-light/10-h-dark photoperiod.

### Observe *F. oxysporum* cells within plant tissue

The staining process using wheat germ agglutinin (WGA) coupled to the fluorophore Alexa Fluor 488 (WGA-Alexa Fluor 488) and Propidium Iodide (PI) is employed for observing fungal colonization in plant leaves and roots following the staining method described previously (46, 81). The infected roots were observed under an Olympus FluoView FV1000 confocal microscope (Tokyo, JP) and photographed with a Hamamatsu camera (Hamamatsu, JP). The experiments were repeated twice.

### Detect *F. oxysporum* from above-ground tissues

To detect Fol4287 and MRL8996 from above-ground tissues, tomato stems were harvested after 16 dpi and washed with abundant SDW. Under the laminar flow hood, stems were dipped in 10% NaClO for 1 min, washed several times with SDW, and blotted dry with Kimwipe paper towels. Small stem pieces were placed on a PDA medium and incubated at 28°C. After 3 days, the observed fungal growth was transferred to new Petri dishes with PDA to obtain axenic cultures. The Qiagen plant DNA extraction kit (Hilden, Germany) was used to extract the DNA of three different fungal colonies isolated from the tomato stems. Fol4287 strain-specific genes FOXG_22560 and FOXG_18682 were amplificated by PCR using the primers: Fol4287_22560_For: ATGCGCTTC AATGTTCTCGC and Fol4287_22560_Rev: ACAACAGACAGTACCAGCGG, as well as Fol4287_18682_For: GCTGCTACGGCGATACTGTC and Fol4287_18682_Rev: GACTCGTCTGGGCTGTACTC with 63 and 64 °C of alignment temperature respectively, 35 cycles and following the manufacturer instructions of Phusion High–Fidelity DNA Polymerase (Thermo Scientific). Finally, the PCR products were visualized in 1.2% of agarose gels.

### Contour-clamped homogeneous electric field (CHEF) electrophoresis

The harvested conidia sample was washed with sterile water, followed by 1.2 M KCl, resuspended in protoplasting solution (25 mg/mL driselase, 5 mg/mL lysing enzyme from Sigma-Aldrich, and 1.2 M KCl), and incubated overnight at 30°C under shaking at 80 rpm. The protoplasts were centrifuged at 4°C for 15 min at 1,000 g, washed by slowly resuspending in 10 mL STC buffer (1 M sorbitol, 10 mM Tris-HCl pH 7.4, and 10 mM CaCl_2_), and centrifuged again as above. The supernatant was carefully poured out, and the protoplasts were resuspended in STE buffer (2% [w/v] SDS, 0.5 M Tris-HCl pH 8.1, and 10 mM EDTA) to a titer of 2 × 10^8^ conidia/mL. The protoplasts were incubated at 50°C for 10 min and mixed at a 1:1 (v/v) ratio with 1.2% (w/v) Bio-Rad Certified Low Melt Agarose. The mixture was transferred to CHEF Disposable Plug Molds and stored at 4°C for long-term storage. CHEF gel electrophoresis was performed as previously described (82). Briefly, the CHEF Mapper System (Bio-Rad) with 1% (w/v) SeaKem® Gold Agarose (Lonza) in 0.5× Tris borate EDTA (TBE) buffer was used to separate chromosomes at 4–7°C for 260 h. The switching time was 1,200 to 4,800 s at 1.5 V/cm, and the angle was 120°. The running buffer was changed every 2–3 days. The gels were stained with 3× GelGreen (Biotium).

### Genome alignment

The alignments of the Fol4287 and MRL8996 nuclear and mitochondrial genomes with repeats masked were generated using MUMmer version 3.23 (83) with the option ‘nucmer--maxmatch’. Alignments less than 1 kb were removed using ‘delta-filter -g −l 1000’. The extent of sequence identity was calculated for each chromosome by multiplying the percentage sequence identity for each alignment by the length of the alignment averaged over the total length of alignments for the chromosome. The circular plot was generated using Circos (84). Only alignments longer than 5 kb and contigs longer than 5 kb were considered. Contigs from the same chromosome were concatenated. A sequence alignment plot for the mitochondrial genome was generated via a PlotMUMmerAlignments.m script (available at https://github.com/d-ayhan/tools). The annotation of the Fol4287 genome was obtained from the 2010 assembly, while the MRL8996 genome was annotated *de novo* at JGI (85, 86). The genes in ACs were identified and their enriched Gene Ontology (GO) terms were analyzed. For Fol4287 genes, locus tag IDs starting with ‘FOXG_’ were used, while for MRL8996, the ‘protein_id’ numbers assigned by JGI were used.

### Repeat analysis

Repeats were *de novo* identified and classified using RepeatModeler v1.0.11 (87). A previously curated library of 69 transposable elements (TEs) was included in the repeat sequence database. Usearch with the option ‘-id 0.75’ was used to cluster the sequences around centroid repeat sequences, and the annotations were fixed manually (88). RepeatMasker version 4.0.7 was used to mask and annotate the genome assemblies (89). TECNEstimator, a bioinformatics pipeline used to quantify the copy numbers of repeat sequences using short reads from whole-genome sequencing data (SRA accessions: SRP140697 and SRP214968), was used to estimate read counts (available at https://github.com/d-ayhan/TECNEstimator). The counts were normalized to the median read coverages of the samples. For both genomes, only one representative sequence in a cluster with the highest copy number was selected for downstream analysis.

### GO term enrichment analysis

The genes harbored by ACs were identified as those located in AC contigs of MRL8996 and AC and unmapped contigs of Fol4287. The orthologous genes were identified using OrthoFinder version 2.3.3 with default options (90). GO terms for the Fol4287 and MRL8996 genomes were downloaded from JGI Mycocosm (JGI-specific genome identifiers: Fusox2 and FoxMRL8996, respectively) (85). The GO_term_enrichment_analysis.m script (available at https://github.com/d-ayhan/tools), which utilizes the hypergeometric cumulative distribution function, was used to analyze the enriched GO terms in ACs. The proteins encoded by the genes included in each term were subjected to conserved domain analysis using an InterPro-based annotated file provided by JGI and a NCBI conserved domain database CD-Search (https://www.ncbi.nlm.nih.gov/Structure/cdd/wrpsb.cgi).

### Phylogenetic analysis

The protein sequence alignments were generated by MEGA11 (91) using the ClustalW algorithm. The phylogenetic tree was reconstructed using FastTree (92) with default options and visualized in iTOL (93).

## Supporting information

Fol4287_TEs

MRL8996_TEs

## Acknowledgements

The authors thank the University of Massachusetts High-Performance Computing Center for providing high-performance computing capacities and assisting in data analysis; we also thank Chris Joyner and David O’Neil of the College of Natural Sciences Greenhouse at the University of Massachusetts, Amherst, for continued support with plant propagation. This project was supported by the National Eye Institute of the National Institutes of Health (R01EY030150) and the Natural Science Foundation (IOS-165241). L.-J.M. is also supported by an Investigator Award in Infectious Diseases and Pathogenesis by the Burroughs Wellcome Fund BWF-1014893; SM is also supported by a Vaadia-BARD Postdoctoral Fellowship. The work (proposal: 10.46936/10.25585/60001360) conducted by the U.S. Department of Energy Joint Genome Institute (https://ror.org/04xm1d337), a DOE Office of Science User Facility, is supported by the Office of Science of the U.S. Department of Energy operated under Contract No. DE-AC02-05CH11231. The funding bodies played no role in the design of the study, the collection, analysis, and interpretation of the data, or the writing of the manuscript.

## Author contributions

L.-J.M., D.H.A., and E.P. designed this study. D.H.A. and L.-J.M. wrote the manuscript with input from all authors. D.H.A., S.M., V.S. S.H, and S.W. performed data analysis. D.H.A., S.A., D.M.S, K.R., S.K., M.E.M, M.C.R, S.Y, and R.R.V. performed experiments. T.A, I.V.G, N.S, E.P and L.-J.M. provided funding support for all experiments and analyses.

## Notes

### Competing Interest Statement

The authors have declared no competing interest.

### Summary of Updates

This revision includes more replicates for both in vitro and in vivo assays and further solidifies the conclusions.

