## Supplementary Figures for "Virulence of *Fusarium oxysporum* strains causing corneal or plant disease is associated with their distinct accessory chromosomes"

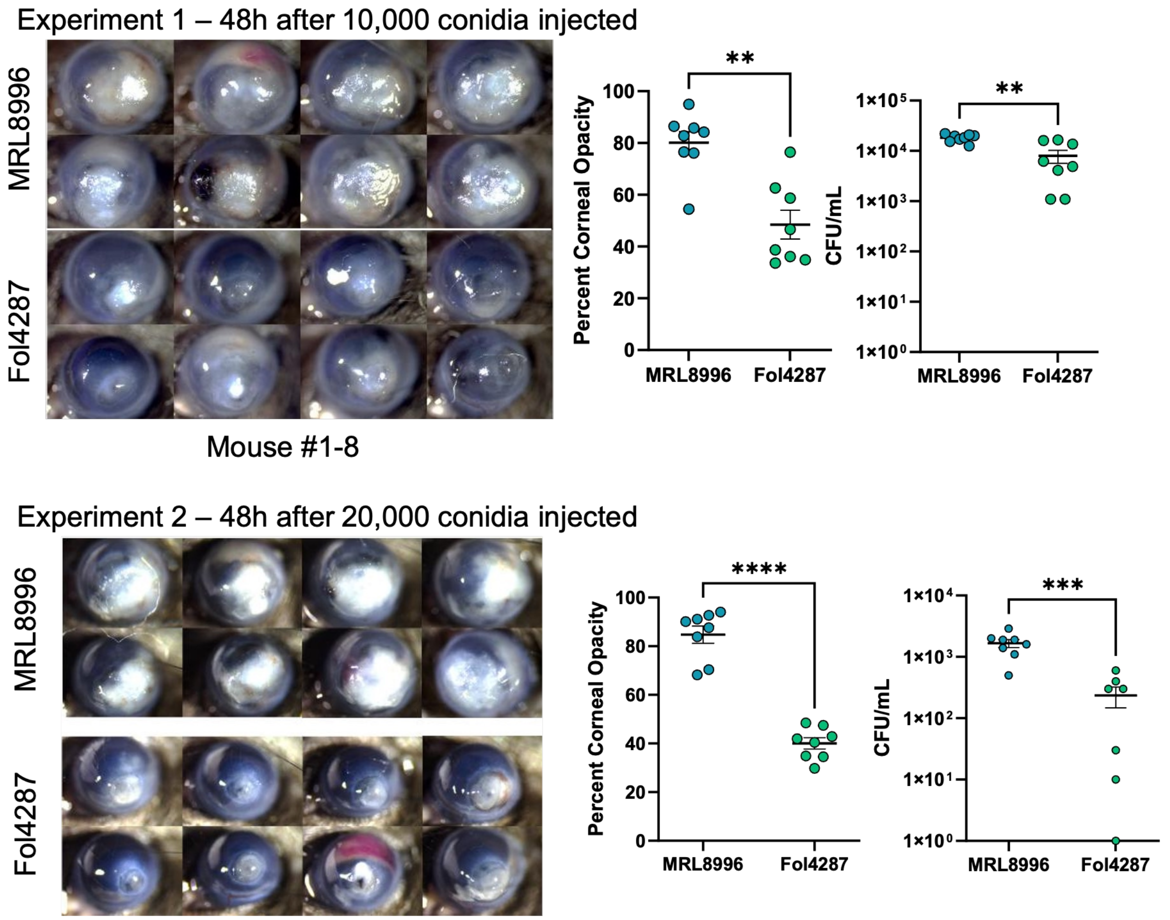


**Figure S1. Additional corneal infection experiment results as described in Figure 1.**

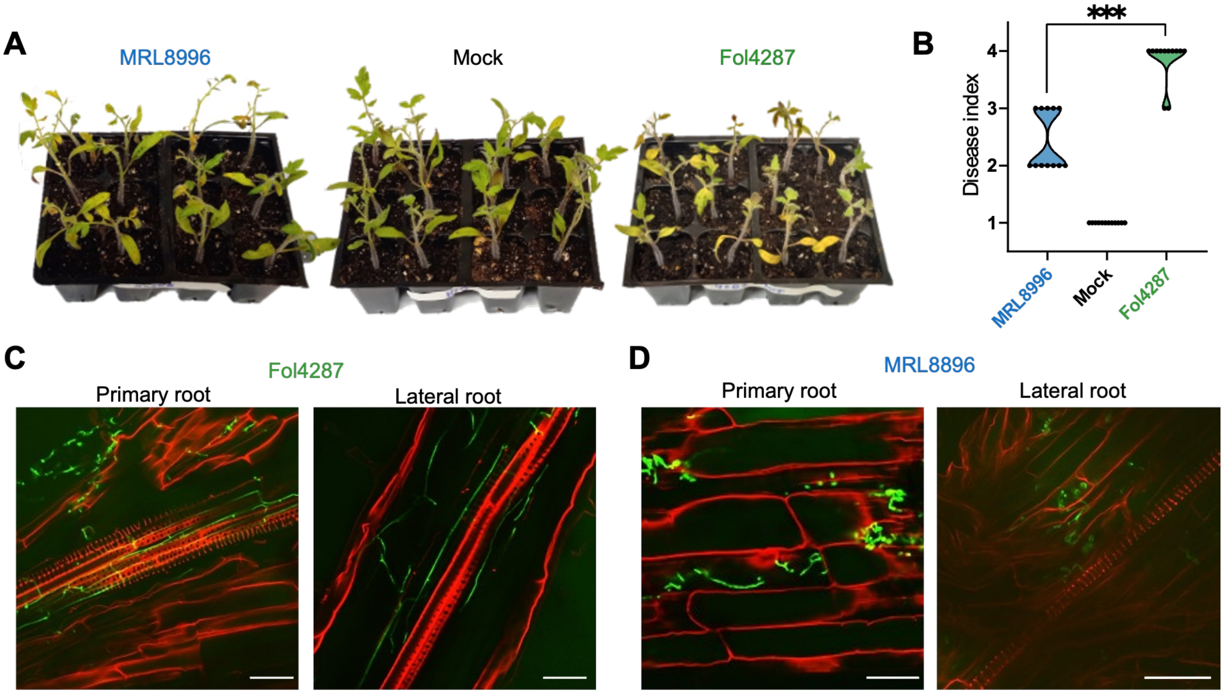


**Figure S2. Additional plant infection experiment results as described in Figure 2.**

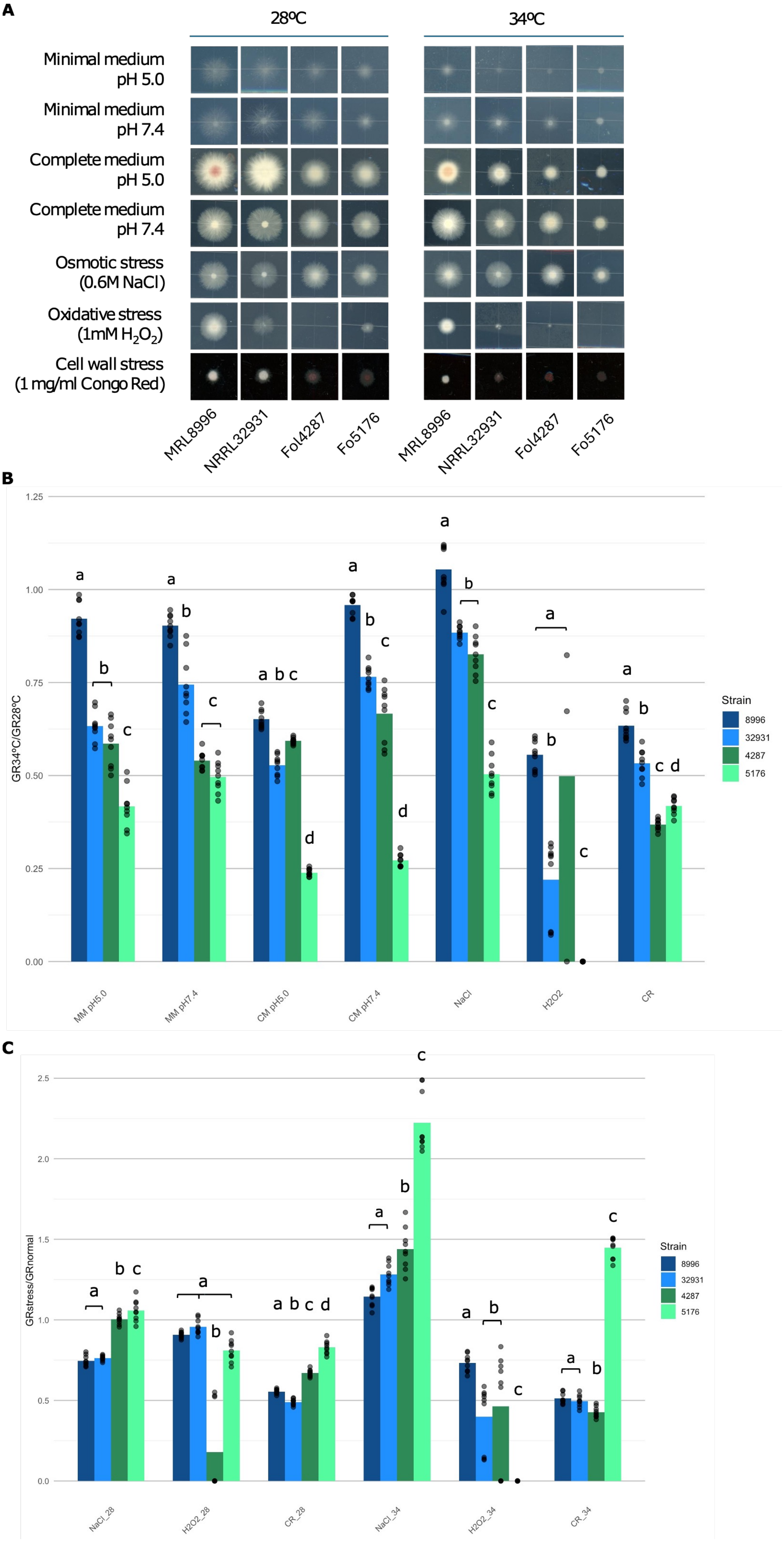


**Figure S3. (A)** ***In vitro*** **growth assays of MRL8996, NRRL 32931, Fol4287, and Fo5276 strains under different abiotic stress conditions. (B) The quantification of temperature tolerance as described in Figure 3B. (C) The quantification of stress tolerance as described in Figure 3C.**

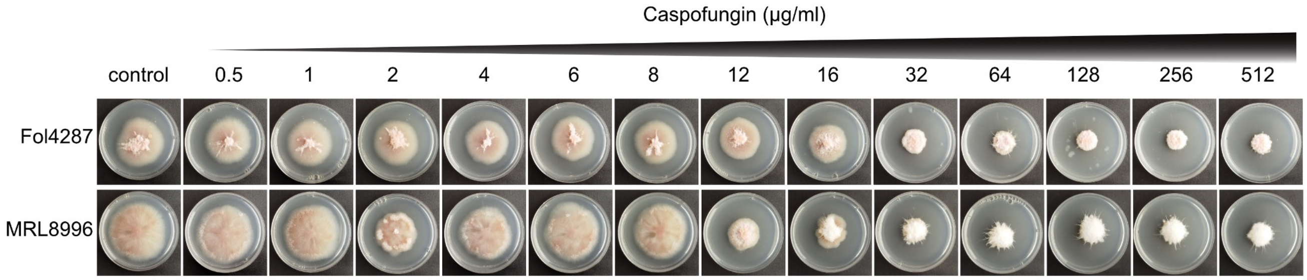


**Figure S4. The colony growth of Fol4287 and MRL8996 under 0 to 512 μg/ml caspofungin concentration in minimal media plates.** Both keratitis (MRL8998) and plant (Fol4287) strains demonstrated enhanced tolerance to high concentrations of caspofungin. Notably, the plant strain exhibited high resistance to caspofungin at concentrations up to 16 μg/mL. In contrast, the growth rate of the keratitis strain MRL8996 dropped by 50% under 2 μg/mL of caspofungin, then reached a 35% decrease at 8 μg/mL, relative to the control (Fig. 4B). This paradoxical caspofungin effect was also observed when measuring the time required to reach 50% conidial germination (Fig. 4C).

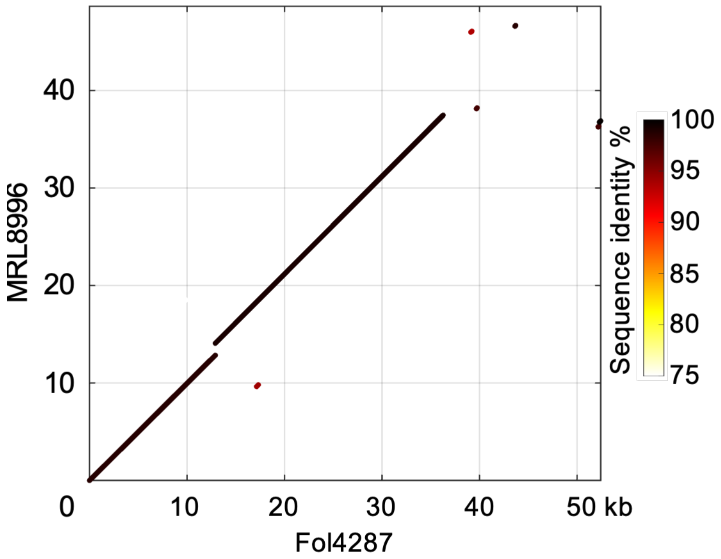


**Figure S5. Dotplot showing the sequence identity between the mitochondrial genomes of Fol4287 and MRL8996.**

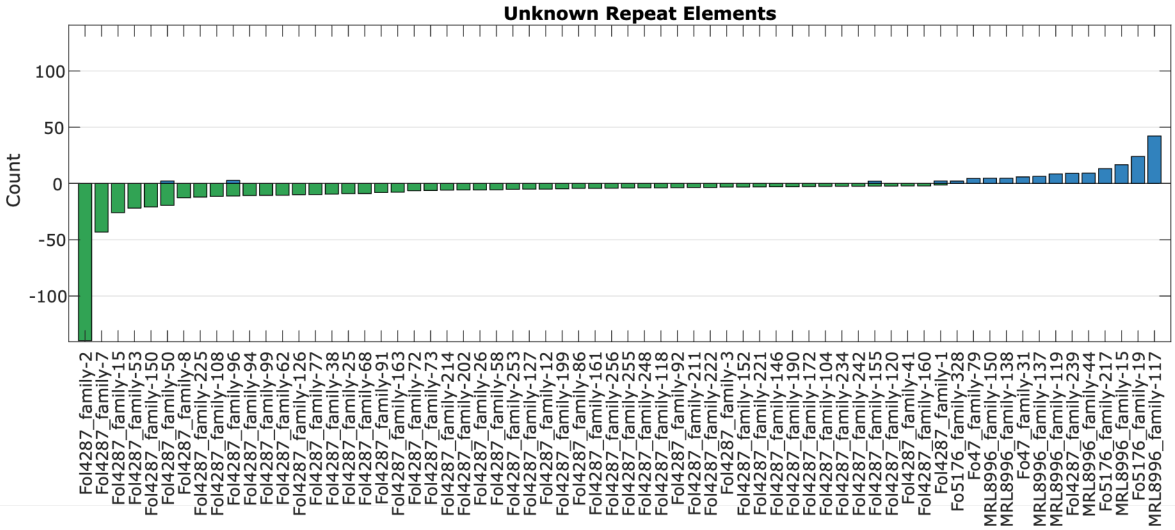


**Figure S6. Counts of unknown repeat families in the Fol4287 (green) and MRL8996 (blue) genomes.**

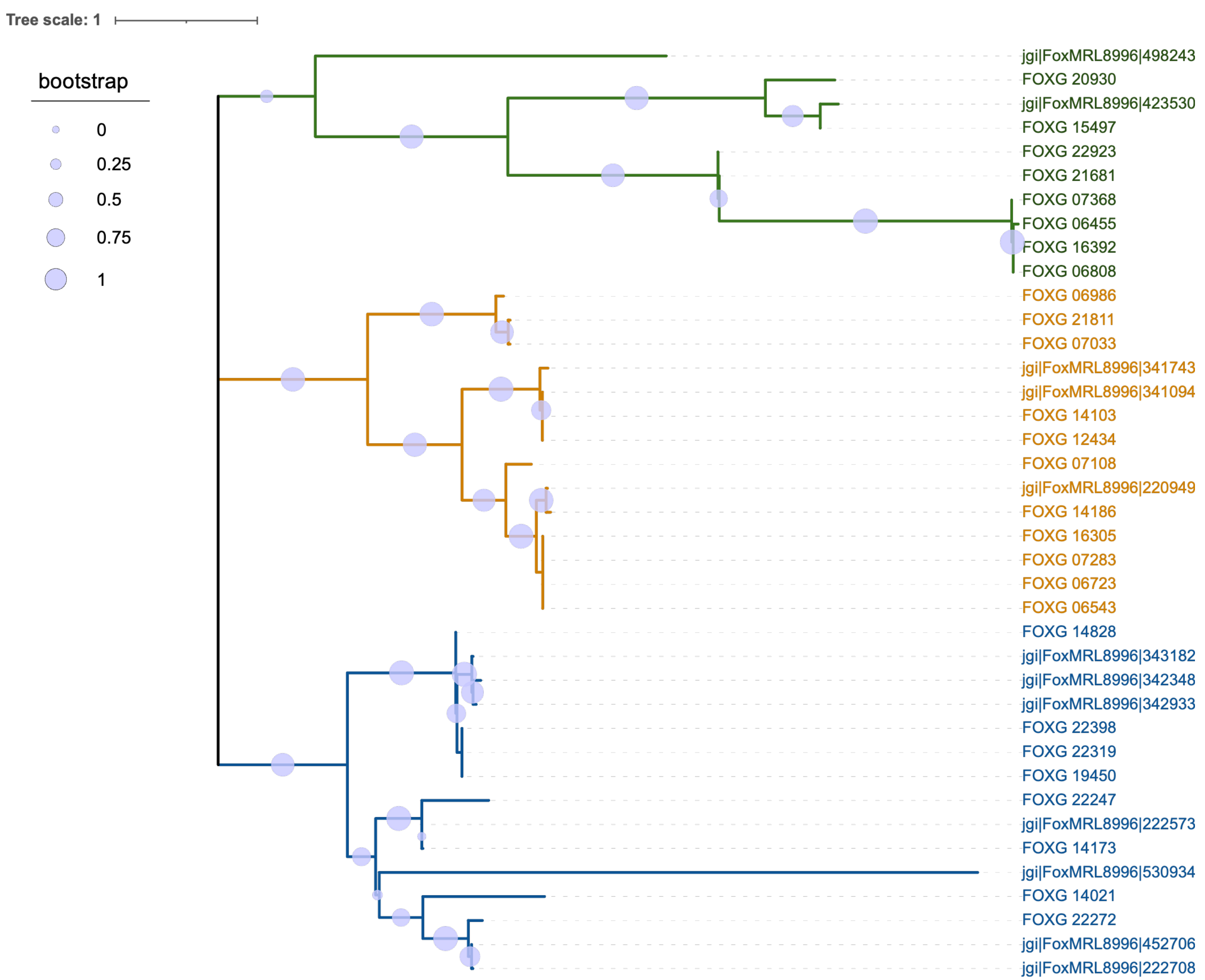


**Figure S7. Phylogeny of proteins involved in chromatin assembly and disassembly encoded by genes located on the accessory chromosomes of Fol4287 and MRL8996.**
